# Adult killifish exposure to crude oil perturbs embryonic gene expression and larval morphology in first- and second-generation offspring

**DOI:** 10.1101/2025.02.10.637481

**Authors:** Jane Park, Chelsea Hess, Charles Brown, David Rocke, Christoph Aeppli, Fernando Galvez, Andrew Whitehead

**Affiliations:** Department of Environmental Toxicology, University of California Davis, Davis, CA 95616, United States; Department of Biological Sciences, Louisiana State University, Baton Rouge, LA 70803, United States; Department of Biomedical Engineering, University of California Davis, Davis, CA 95616, United States; Bigelow Laboratory for Ocean Sciences, East Boothbay, Maine 04544, United States

**Keywords:** Trans-generational, multi-generational, oil spill, developmental toxicology, transcriptomics

## Abstract

Exposures to environmental toxicants can have both immediate and long-term impacts, including those that persist into the next generation. Using Gulf killifish (*Fundulus grandis*), we tested whether adult progenitor exposure to crude oil caused perturbations to larval morphology and embryonic genome-wide gene expression in their first- and second-generation descendants raised in clean water. We also tracked responses of additional direct oil exposures in the F1 and F2 embryos. Exposure to oil in progenitor fish caused altered larval morphology in F1 and F2 descendants. Some perturbations were enhanced by additional oil exposure in lineages with progenitors that had been exposed to oil. Progenitor exposures altered embryonic gene expression in F1 and F2 descendants, implicating impacts on neurological and cardiovascular systems. Molecular responses to progenitor exposure were distinct between F1 and F2 offspring, suggesting complex interactions between mechanisms that contribute to the transgenerational transfer of information. In contrast, molecular responses to additional direct oil exposures during early-life development were highly conserved between generations and between progenitor exposure treatments. We conclude that exposure to crude oil causes developmental perturbations that propagate at least two generations, and we present additional hypotheses about underlying molecular mechanisms and emergent health consequences.

**SYNOPSIS:** This study shows that exposure of adult fish to crude oil perturbs development and gene expression in offspring and grand-offspring, thereby extending the timeframe necessary to consider ecotoxicological risk.

## INTRODUCTION

Responses to environmental stressors can manifest and persist across several biological timescales. Within the lifetime of an organism, environmental perturbations can cause acute or chronic physiological changes. Environmental changes that affect organismal fitness and that persist across many generations may lead to population demographic changes such as population decline or evolutionary adaptation. Effects of environmental exposures can also propagate across intermediate timescales, such as across one or a few generations (transgenerational or multigenerational effects). Environmental information may transfer from parent to offspring and beyond through non-genetic modes of inheritance such as maternal provisioning or epigenetic imprinting ^1–3^. Compared to responses that emerge at physiological or evolutionary timescales, responses that emerge across these intermediate transgenerational timescales are less understood. Important and outstanding questions include: What kinds of environmental changes may have effects that propagate across generations? Are there features of environmental perturbations that allow prediction of whether transgenerational effects are adaptive, neutral, or deleterious? What mechanisms govern the transfer of environmental information across generations? What are mechanistic features that contribute to whether transgenerational impacts are adaptive or deleterious? Progress in answering these questions will be important for improving our ability to assess ecological risks for populations experiencing rapid environmental change in the Anthropocene.

Transgenerational effects may result in health outcomes that are beneficial, neutral, or harmful (e.g., adaptive, neutral, or maladaptive) (Figure 1). These alternate outcomes likely depend on whether the stressor is novel or commonly encountered within a species’ natural niche (evolutionary history) ^4,5^, the nature of exposure such as severity and duration ^6,7^, and whether the environmental change is transient or persistent across generations ^8^. For *adaptive* transgenerational plasticity, resilience to stressors is increased in the offspring of parents who were exposed to those stressors compared to the offspring of parents who were not exposed (Figure 1, blue line). In some cases, this increased resilience in the perturbed environment may come at the cost of reduced resilience in the non-perturbed environment (Figure 1, green line). In adaptive scenarios, the stressor is usually one that is normally encountered within that species’ natural niche, such that the mechanism that transmits environmental information to offspring presumably evolved and is maintained through natural selection. Examples include *Daphnia* exposure to predators triggering the development of increased helmet size in their offspring ^9^ and thermal exposure of sheepshead minnows triggering increased growth rate in their offspring ^10^.

**Figure 1:**
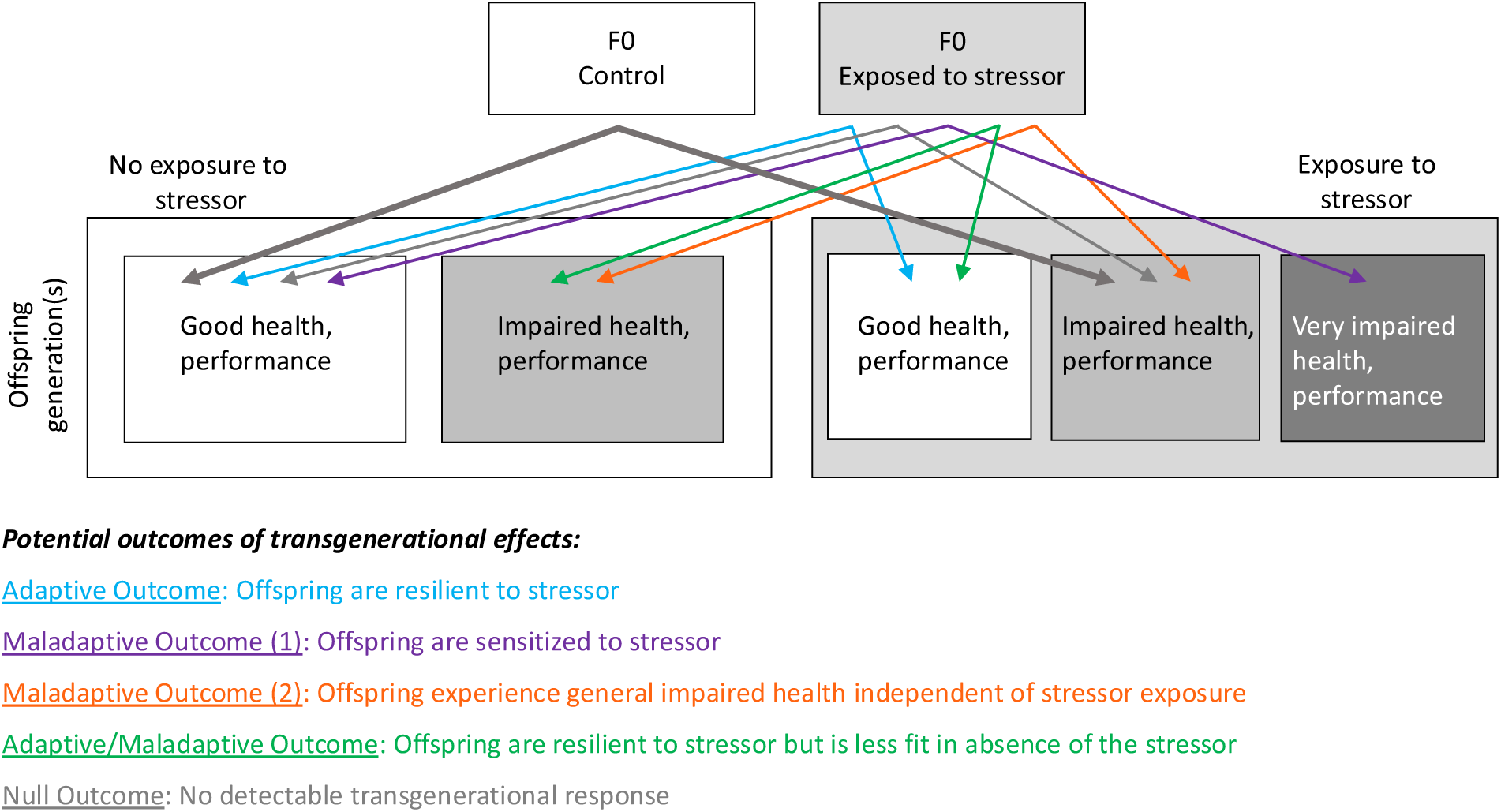
Environmental stressors in the progenitor (F0) generation can cause heritable changes to health or performance of F1 or F2 offspring. These changes can be beneficial (adaptive) or harmful (maladaptive), and likely depend on the offspring environment and ecological familiarity of the stressor to the organism. Transgenerational effects that alter responses to the specific stressor can lead to resilience (blue arrows) or sensitivity (purple arrows) to the stressor in F1 or F2. Stressor-specific transgenerational effects can have fitness tradeoffs if F0 and F1 environments differ (green arrows). Transgenerational effects can also cause general impairments of health that are non-specific to the stressor (orange arrows).

For agents of environmental changes that have no analogue in that species’ ecological or evolutionary history, negative or *maladaptive* transgenerational outcomes may be more likely. In these cases, exposures that cause health impairments may have negative consequences for offspring, for example through impaired egg provisioning or parent-to-gamete carry-over of small molecules (e.g., chemical toxicants) that perturb development. Furthermore, no-analog environments may cause malfunctioning of the molecular machinery that transmits cellular memory such as epigenetic imprinting systems ^11^, such that developmental programs and subsequent health are perturbed. This could result in increased sensitivity to the original stressor (Figure 1, *purple* line); but since the molecular malfunctions may be non-specific, the health impairments could also manifest as non-specific maladies that are difficult to predict or anticipate. Furthermore, these maladies may manifest in otherwise benign offspring environments (Figure 1, *orange* line). In the Anthropocene, no-analog environments (pollution, climate change, invasive species) are likely emerging too swiftly and severely for adaptive trans-generational mechanisms to evolve in most species. For them, perturbations of transgenerational mechanisms may contribute to health impacts thereby amplifying population decline. Therefore, characterizing these impacts, and their underlying mechanisms, is important for risk estimation and assessment. We consider how toxicant exposures may have impacts that propagate across one or two generations, thereby revealing the influence of mechanisms including maternal carry-over effects and those that are likely to occur through epigenetic alterations.

Chemical toxicants are an important component of the Anthropocene, but risk estimation and assessment of chemicals typically do not consider transgenerational effects ^12^. One might imagine several different transgenerational outcomes following toxicant exposure. Offspring could be more resilient to a toxicant because maternal carry-over of a chemical to eggs may trigger molecular programs in developing embryos that are protective in the newly polluted environment (e.g., detoxification pathways). This effect would persist beyond one generation only if exposures also persist. Alternatively, offspring may have impaired health following parental exposures. This could result from exposures that reduce egg or sperm quality, impair energetic provisioning, or that cause chemical carry-over into eggs thereby perturbing development. These outcomes would persist beyond one generation only if exposures also persist. In contrast to these outcomes, exposure effects that propagate more than one generation - in the absence of subsequent exposures - would indicate perturbations of cellular memory (e.g., epigenetic changes) thereby altering development with negative consequences for health, performance, and fitness. We predict that toxicant perturbations of cellular memory are unlikely to be beneficial because responses to no-analog stressors are unlikely to have been shaped and canalized by natural selection. In this case, indiscriminate perturbation of cellular memory machinery may manifest in subsequent generations as phenotypic alterations that are complex, unpredictable, and maladaptive.

We integrated transcriptomics with measures of developmental morphology that are often perturbed by oil exposure to track impacts of crude oil exposures across two generations in Gulf killifish (*Fundulus grandis*), an ecologically important species that is abundant in habitats impacted by the Deepwater Horizon oil spill ^13,14^. Oil spills are a notorious feature of the Anthropocene ^15^. Oil exposures are often discontinuous, at least for relatively long-lived species and those with medium to large home ranges. Exposures cause a diverse array of acute and chronic health impairments ^16–18^, but whether exposures cause impacts that propagate to future generations is not well characterized. Some recent studies in invertebrates have documented crude oil exposures that cause multigenerational impacts ^19–22^, but few have examined effects that propagate across more than one generation ^20,22^, and the same appears true for fish ^23–26^. Studies that examine oil effects in fishes that propagate across more than one generation are few (e.g., ^27^). Moving forward, it is important to understand whether the effects of crude oil exposure propagate across one or more generations and the array of underlying molecular mechanisms and physiological consequences in at-risk species.

Following exposure to sub-lethal levels of crude oil for 36 to 48 days, adult *F. grandis* were spawned. Their offspring are considered members of the “progenitor oil exposed lineages”. We generated a second set of control lineages whose progenitors had not been exposed to oil, where their offspring are considered members of the “progenitor no-oil control lineages”. We tracked descendants of these two sets of lineage progenitors (F0) for one and two generations (F1 and F2, respectively). In descendent F1 and F2 generations, we tested whether progenitor exposure, with and without additional direct embryonic exposures to crude oil, affected 1) larval morphology, and 2) genome-wide gene expression throughout embryonic development. We considered two alternate hypotheses regarding the transgenerational effects of crude oil exposure: 1) Progenitor (F0) oil exposure causes heritable perturbations of development in F1 and F2 descendants, and 2) Progenitor oil exposure enhances offspring sensitivity to additional direct oil exposure. Since natural populations do not normally encounter crude oil, we predicted that transgenerational impacts, if any, would be negative. It is important to note that if transgenerational impacts are indiscriminate, phenotypic outcomes could be unpredictable. We, therefore, tracked genome-wide gene expression, which, if perturbed by progenitor exposure, could stimulate hypotheses about the nature of emergent phenotypic outcomes.

## MATERIALS AND METHODS

### Exposure of F0 adults to the water-accommodated fraction (WAF) of crude oil

All procedures that are described were performed in accordance with protocols approved by the Institutional Animal Care and Use Committee at Louisiana State University (Protocol: 15-070). Adult *F. grandis* originally from Leeville, LA (29°15’24.6“N 90°12’51.3” W) were held in laboratory conditions at Louisiana State University for at least 2 generations before experiments. Two separate parental oil exposures were performed. In the first exposure, adult F0 (progenitor) fish were exposed for 36-45 days to either control water (*n*=19 fish) or weathered crude oil (*n*=26 fish) at a salinity of 12 g/L made with artificial Instant Ocean at a natural light cycle. Each treatment was conducted in four 70-L glass aquaria per treatment at a density of 5-7 adult *F. grandis* per tank. These progenitors were used to generate the F1 generation fish, which were grown into F1 brood stocks to generate F2 generations (see below). Fish used in the F1 broodstock were never directly exposed to oil. Polycyclic aromatic compound (PAC) analyses of water samples from these exposures were performed on 150 mL composite water samples collected daily (50 mL per tank from 3 random tanks) for 6 to 7 days. Each composite WAF sample (*n*=6) was acidified with organic-free hydrochloric acid at a ratio of 2 mL acid to 1 L of water to a pH < 2 and held at 4 °C awaiting extraction (USEPA method 3510C). Samples were shipped to ALS Environmental (Kelso, WA) for PAC analysis and alkylated homologs using GS-MS (USEPA method 8270D). The mean PAC concentration was 76.0 ± 42.5 µg L^-1^ based on the cumulative concentrations of a total of 39 PACs (See Appendix S1 for the list of 39 PACs). In the second exposure, adult F0 (progenitor) fish were exposed for 40-48 days to either control water (n= 32 fish) or weathered crude oil (n=37 fish) using similar conditions as described for the first exposure. These fish were used only as brood stock to obtain F1 generation embryos for acute toxicity testing and morphological assessment of F1 embryos during embryonic development. This second exposure was necessary since all of the F1 individuals from the first exposure were needed to serve as broodstock to obtain an F2 generation that excluded pairing of close relatives. Both exposures were conducted using similar procedures. The mean PAC concentration of the second F0 exposure was 119.1 ± 28.4 µg L^-1^ based on the cumulative concentrations of the same suite of 39 PACs as in the first exposure.

Because of the large volume of WAF needed, the traditional method of producing WAF by high-energy mixing was not feasible. As such, a high-volume WAF generator was constructed consisting of a 1000-liter recirculating system lined with a polytetrafluoroethylene (PTFE), a water pump at the base, and PTFE pipe that recirculated water from the bottom to the surface. Water was passed through oiled sintered glass beads (Siporax®, 15 mm, Sera) in a PTFE-mesh bag contained within the upweller pipes as described by Carls et al. ^28^ and by Kennedy and Farrell ^29^. The beads were pretreated with Macondo-252 surrogate light sweet crude oil (supplied by BP America Production Company; sample ID: SO-20110802-MPDF-01) at a ratio of 3.74 g oil per g bead for 48 h at 4 °C in a 2.5 L amber bottle. Oiled beads were poured into a narrow-stem glass funnel to remove the non-adsorbed oil from the beads and then loaded to a chamber in the upweller pipe at a ratio of 0.67 g bead L^-1^ artificial seawater (ASW) (Instant Ocean salt mix; United Pet Group, Cleveland, OH, USA). This represented an oil loading rate of 2.5 g L^-1^ water, which was similar to that described by Pilcher *et al*. ^30^ during adult oil exposures to Macondo oil. Beads soaked in water served as the control. Water was passed across the Siporax beads for 24 hours. After 24 h, the pump was turned off, allowing an oil sheen to form. The water-accommodated fraction was drained from the bottom to prevent the collection of the oil sheen. On water change days, after the control and WAF were used for tank water replacements, the 1000-L tanks were refilled with 12 g/L artificial salt water, and the process was repeated to generate new batches of water as described above. For exposures, water from WAF tanks and clean tanks was transferred to fish exposure tanks. Every four days, ∼800 L of clean and WAF treatment waters were produced to provide 50% water exchange (30 L every 4 days) for thirty 60-L tanks (15 tanks for controls and 15 tanks for WAF exposures).

### Generation of F1 and F2 embryos for transgenerational experiments

Following exposure to oil or control conditions, adult fish were used as broodstock to collect gametes for the generation of F1 embryos (Figure 2). We created two lineages – one where progenitor males and females were exposed to oil (progenitor “exposed” lineage), and a control lineage where progenitors were not exposed to oil (progenitor “control” lineage). Eggs were stripped from individual control and oil-exposed females and combined with sperm from individual males from the same treatment, followed by adding 12 g/L water to activate gametes. This procedure was repeated until at least four to five unique female-male pairings were created for each lineage. These unique female-male pairings are subsequently referred to as families. Two separate spawning events took place on experimental days 36 and 45 to generate enough families per lineage. By day 45 mortality in the oil-exposed adults was low (∼12.8%). The two lineages were made using the following combination of gametes: control female x control male, and oil-exposed female x oil-exposed male. Embryos were assessed for successful fertilization 1 hour later (hardened chorion layer and raised fertilization envelope). Unfertilized embryos were removed. Families were reared separately until large enough (∼4 months of age) before they could be tagged with unique combinations of visible implant elastomer (VIE) tags below the dorsal fin (Northwest Marine Technology, Inc.) to identify the progenitor history and family. After tagging, families were combined to facilitate holding under common-garden conditions until F1 fish were sexually mature. Tagging was repeated as needed. For generating F2 offspring, multiple independent crosses were performed to avoid full-sibling mating of F1 individuals. F1 and F2 offspring from control and oil-exposed F0 progentiors were reared in clean conditions (12 g/L ASW) throughout their lives. A subset of F1 and F2 embryos was tested for their sensitivity to additional direct oil exposure (Figure 2).

**Figure 2:**
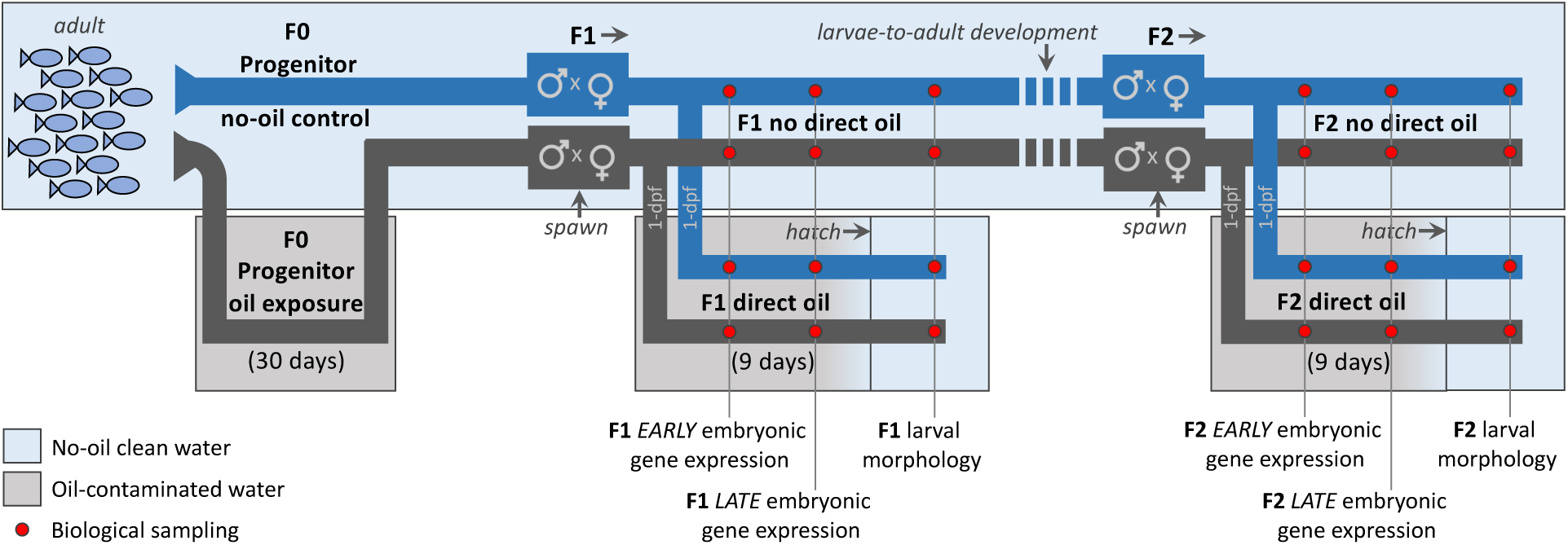
Lineages of no-oil control (blue line) or oil-exposed (gray line) F0 progenitor adults were established. Fish were mostly housed in clean water (blue boxes) except for limited exposures to oil-contaminated water (gray boxes). First- and second-generation (F1 and F2) descendants were created by mating males and females within each lineage. F1 adults mated to create the F2 generation had not been directly exposed to oil during their lifetime. Subsets of F1 and F2 descendants were directly exposed to oil during embryogenesis starting at 1-day post-fertilization (1 dpf) until hatch. Biological samples (red dots) were collected for genome-wide gene expression analysis and larval morphometrics. Biological samples were collected to contrast responses to two main effects: that of direct embryonic oil exposure (gray water vs. blue water) and that of progenitor oil exposure history (blue line vs. gray line).

### Generation of F1 embryos for acute toxicity testing and morphological assessment

Eggs were stripped from individual females drawn from control and oil-exposed conditions and combined with sperm from males drawn from the same exposure condition, followed by adding 12 g/L water to activate gametes. These F1 embryos were then directly exposed to acute oiling and control conditions as described below. Similarly, the F2 embryos derived from the F1 adult fish were used in direct exposures to acute oil. These early-life F1 and F2 embryos were assessed for mortality, morphological impairments, and gene transcription.

### F1 and F2 embryonic direct oil exposure

A subset of embryos was exposed directly to sub-lethal levels of a high energy water accommodated fraction (HEWAF) of oil until hatch for up to a 21-day period. HEWAF was generated based on the protocol described in Incardona *et al*. ^31^, where artificial sea water (ASW) was created by mixing artificial sea salt (Instant Ocean salt mix; United Pet Group, Cleveland, OH, USA) in reverse osmosis-filtered water to a salinity of 12 g/L. An initial loading concentration of 2 g of surrogate Macondo oil per liter of ASW was used to generate HEWAF preparations. HEWAF was generated by vortex (15,000 RPM) in a Waring CB15 blender (Waring; Torrington, CT, USA) for 30 seconds. This mixture was transferred to separatory funnels to settle for 1 hour before the bottom HEWAF fraction was drained into a glass beaker. Care was taken to avoid the collection of any slick residue that had formed at the top of the mixture. Undiluted HEWAF was then transferred to a 75-L glass aquarium and the process was repeated until a sufficient volume of WAF was produced. The undiluted HEWAF mixture was partitioned between 1-L glass bottles and held at −20 °C until needed, at which point they were thawed overnight while kept in the dark. This common source of 100% HEWAF was used for all embryonic exposures. As needed, HEWAF was diluted serially to 56%, 32%, and 10% of the original HEWAF with 12 g/L ASW, herein referred to as high, medium, and low concentration treatments. All exposure concentrations were sub-lethal. Exposures were performed using 250 mL HEWAF in Pyrex dishes under constant gentle agitation for 21 days. Every two days, the entire volume of water was replaced with freshly prepared HEWAF. For the acute exposures of the F1 embryos, the mean PAC concentrations were 53.3 ([low]), 138.0 ([medium)], and 227.9 µg/L ([high]), and for the F2 embryo exposures, the mean PAC concentrations were 47.7 ([low]), 119.5 ([medium)], and 241.7 µg/L ([high]). Embryos were examined daily for mortality and the presence of specific morphological markers to ensure consistent sampling across developmental time. Embryos were flash frozen in liquid nitrogen at post-neurulation (∼2 days dfp, “early stage”; (Armstrong and Child 1965)) and pre-hatch/onset of eye and pectoral fin movement (∼15-20 dpf, “late stage”); these embryos were preserved for RNA-seq analysis.

### Larval morphology

Morphometric data were collected from F1 and F2 larvae derived from the acute embryonic oil exposure (and control) experiments described above. These fish were reared in glass dishes separate from embryos that were reared for transcriptomic sampling, in the same clean or oiled conditions and using the same batch of prepared HEWAF as those sampled for transcriptomics. Morphometric data were collected from [control], [medium], and [high] direct embryonic oil exposure treatments for F2 fish, but these data were collected from only [control] and [medium] treatments for F1 fish because of limited samples. Upon hatching, larvae were immediately fixed in 4% paraformaldehyde, and images were taken along the sagittal plane using Zeiss SteREO Lumar V.12. Total length was measured from the tip of the head to the end of the caudal fin. The pre-orbital length (POL) was measured from the tip of the head to the eye. All measurements were processed using Zeiss Zen (Blue Edition).

Treatment effect on larval morphometric endpoints were tested using two-factor ANOVA in the R package *car* v3.1-2 ^32^. The statistical model included two main effects: the F0 progenitor oil treatment (control or exposed) and the descendant embryonic direct oil exposure treatment (control or exposed) - and their interaction. Prior to ANOVA data were confirmed to be normally distributed using the Shapiro Wilk test in R. If main effects or interactions were statistically significant (p<0.05) Tukey posthoc tests were performed in the R package *emmeans* v1.9.0 ^33^. All R analyses were conducted using Rstudio v2023.9.0.463 ^34^.

### RNA-Sequencing and read count quantification

Messenger RNA was extracted from flash-frozen whole embryos using Zymo Directzol 96 (Cat # R2057). RNA-Seq libraries were prepared using NEBNext Directional RNA Library Prep Kit for Illumina (Cat # E7420L) which included poly-A selection for mRNA. Libraries were sequenced on HiSeq 4000 as paired end 150-bp reads at the UC Davis Core Genomics Facility. Approximately 12-13 million raw reads were obtained per individual sample, including five replicate individual samples per treatment group. There were eight treatment groups in total per generation: two F0 progenitor exposure lineages (no-oil control and oil-exposed lineages), two embryonic direct-exposure conditions (control and oil-exposed) and two embryonic sampling timepoints (early and late stages of development). Raw reads were checked for quality using FastQC ^35^, trimmed with trimmomatic ^36^, and mapped and quantified using Salmon ^37^. Reads were mapped to the *Fundulus heteroclitus* gene model set annotated for the reference genome by Reid et al. ^38^. Gene-level counts were estimated from transcript-level estimates using tximport ^39^.

### Differential gene expression analysis

Each generation was analyzed independently. This is because library preparation and sequencing for F1 and F2 experiments were conducted in different years, thereby potentially introducing batch effects that could confound direct comparisons of F1 and F2 results. Data from each developmental stage were analyzed independently to simplify our statistical analysis, as the developmental stages that we examined have massive differences in gene expression as observed in other studies that contrast gene expression across developmental stages ^40–42^, and because we were not interested in treatment effects between very early and late embryogenesis. Genes with low read counts were excluded from analyses. Low counts were defined as < 5 raw counts in < 4 replicate embryos in every lineage-by-embryo exposure treatment group, such that genes that are transcriptionally active (>5 counts in at least 4 replicate individuals) in at least one treatment group would be retained. We fit our data to meet the assumptions for linear regression modeling using the methods described in Rocke et al. (2015). Log-transformed counts were normalized to the grand mean. We subtracted the mean log-transformed gene count of each sample, *i*, and added the grand mean of log counts of all genes and samples from every log-transformed count for each gene in sample *i*. Differential gene expression was tested using a standard linear regression model in R ^43^ defining F0 progenitor oil treatment (*X_i_*) and descendant embryonic direct oil exposure treatment (*Z_i_*) as main effects and the interaction effect between the effects (FDR-adjusted p < 0.1).

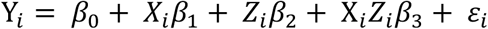

### Gene Ontology (GO) enrichment analysis

Differentially expressed genes (DEGs) for the main and interaction effects were filtered by those with orthologs associated with human and zebrafish Uniprot accession IDs. Hierarchical clusters (Pearson correlation) of DEGs with similar expression patterns were tested for functional annotation enrichment using DAVID Bioinformatics Resources 6.8 ^44^.

## RESULTS

### Larval morphometrics

We detected no elevated mortality in oil exposed embryos compared to controls, confirming that exposures were within the intended sub-lethal range. We observed that direct embryonic oil exposure caused reduced larval total length that was consistent whether or not progenitors had been exposed to oil, and these effects were also consistent in F1 and F2 generation exposures (Figure 3A, 3B, 3C). If progenitors had been exposed, F2 descendants had reduced total length (Figure 3B, 3C); we observed a similar progenitor-effect trend in F1 descendants, but this was not statistically significant (Figure 3A; p=0.09). Progenitor exposures did not modulate the sensitivity of the larval total length response to direct embryonic exposures (no significant interactions between progenitor exposure and embryo direct exposure main effects). In summary of total larval length results, progenitor exposure did not sensitize descendants to direct oil exposure but had a subtle impact on descendant’s length regardless of direct exposure. This is consistent with the *orange* outcome (*Maladaptive Outcome 2*) in Figure 1.

**Figure 3:**
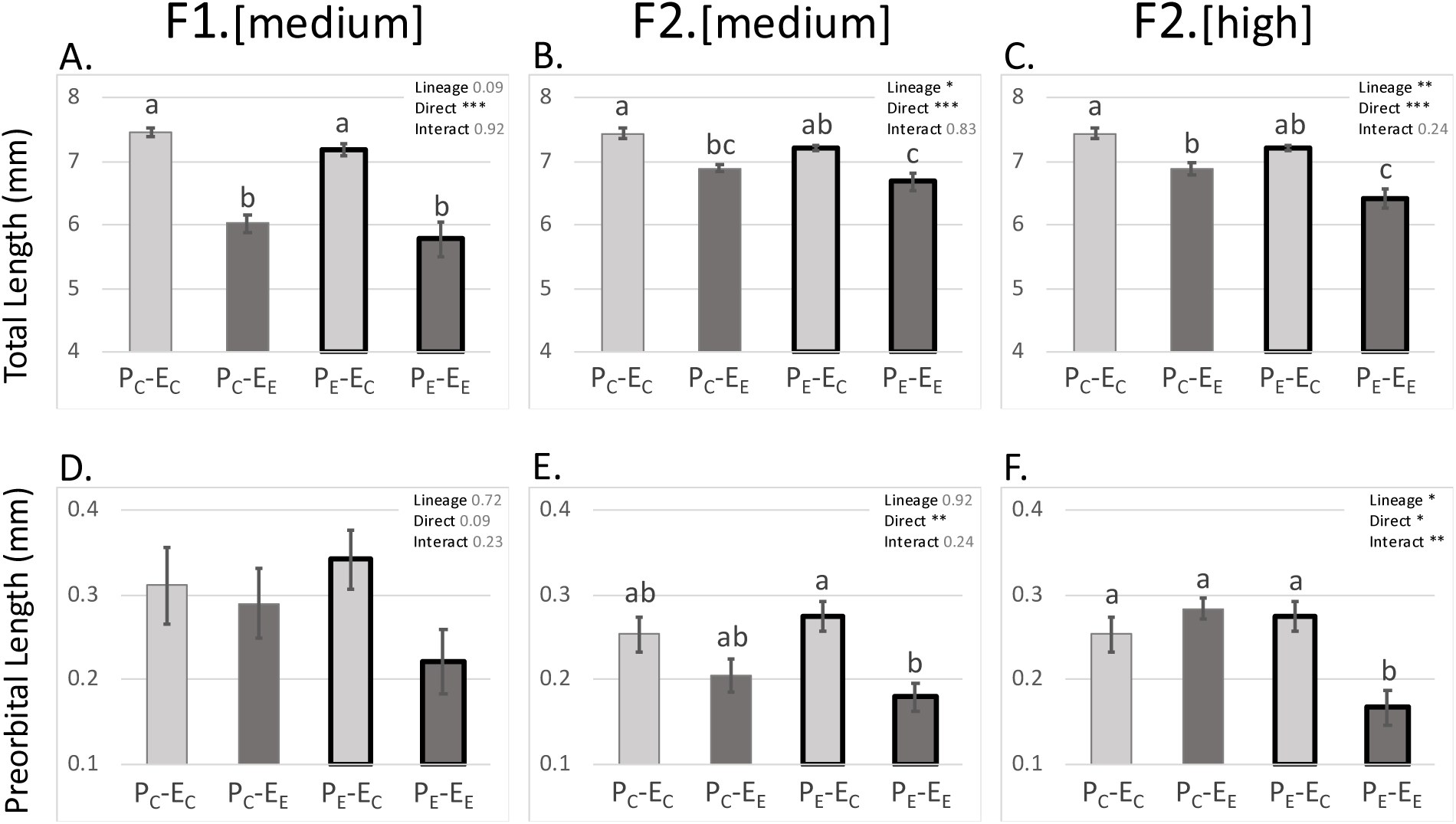
Treatment effects of progenitor and direct-embryonic oil exposures on F1 and F2 larval morphology. Treatments (bar labels) include progenitor oil exposure condition including progenitor no-oil control (P_C_) and progenitor exposed (P_E_), and embryo direct oil exposure condition including embryo no-exposure control (-E_C_) and embryo exposed (-E_E_). Accordingly, the pair of bars on the left side of each panel with the thin gray outline are data from the progenitor no-exposure control conditions (P_C_-E_C_ and P_C_-E_E_), and the pair on the right with the bold dark outline are from the progenitor oil exposed conditions (P_E_-E_C_ and P_E_-E_E_). Within the progenitor exposure condition pairs, the bars with light gray fill indicate the embryo no-oil exposure control conditions (P_C_-E_C_ and P_E_-E_C_), and the bars with the dark gray fill indicate the embryo direct oil exposure condition (P_C_-E_E_ and P_E_-E_E_). Plots show the treatment effects on total larval length (A. through C.) and larval pre-orbital length (B, C, D). Plots are included for F1 (A, D) and F2 (B, C, E, F) descendants. F2 outcomes are shown for embryo medium-dose exposure (B, E) and embryo high-dose exposure (C, F). Within a plot, outcomes (p-values) for statistical tests are shown for the two main effects including the progenitor exposure treatment (lineage) and embryo direct exposure treatment (direct) and their interaction. p<0.05, p<0.01, and p<0.001, are indicated by *, **, and ***, respectively. The same letter above each bar indicates no significant difference between treatment means following post-hoc tests. Error bars represent standard errors of the mean.

We observed that direct embryonic oil exposure caused a reduction in larval pre-orbital length. This response was statistically significant in F2 embryos; a similar response was observed in F1 embryos, but this was not statistically significant (p=0.09). Progenitor oil exposure modulated (enhanced) the sensitivity of the response to direct embryonic exposures in F2 descendants (at the higher dose; significant interaction, p<0.001; Figure 3F). Enhanced sensitivity was suggested by trends in other treatments (e.g., F1) but interactions were not statistically significant. In summary of pre-orbital length results, progenitor exposure sensitized descendants to direct oil exposure in some conditions, which is consistent with the *purple* outcome (*Maladaptive Outcome 1*) in Figure 1.

### Gene expression

Similar numbers of genes were included in our four separate analyses after filtering low-abundance transcripts (F1 early stage: 20,044, F1 late stage: 23,552, F2 early stage: 20,356, F2 late stage: 23,465) (Tables S1 to S4). We detected greater numbers of differentially expressed genes (DEGs) in response to experimental treatments (progenitor exposure lineage, direct embryonic exposure treatment, and their interaction) that included late-stage embryos compared to those that included early-stage embryos (Table 1).

**Table 1:** Numbers of differentially expressed genes for experimental main effects and interactions (rows) for F1 and F2 embryos examined at early and late stages of development.

|  | F1 |  | F2 |  |
| --- | --- | --- | --- | --- |
|  | Early Stage | Late Stage | Early Stage | Late Stage |
| Progenitor Oil Exposure Lineage | 2 | 165 | 81 | 292 |
| Direct Embryonic Oil Exposure | 8 | 639 | 14 | 716 |
| Interaction between Direct Embryonic Exposure and Progenitor Exposure Lineage | 0 | 6 | 4 | 53 |

### Progenitor exposure to oil alters developmental gene expression in descendent F1 and F2 embryos

We observed that progenitor exposure to oil perturbed developmental gene expression in F1 and F2 embryos. This progenitor exposure effect was greater in late-stage development compared to early-stage, and greater in F2 descendants compared to F1. In the F1 generation, progenitor exposure changed the embryonic expression of 2 and 165 genes in early-stage and late-stage embryos, respectively (Table 1). In the F2 generation, progenitor exposure changed the embryonic expression of 81 and 292 genes in early-stage and late-stage embryos, respectively.

### Progenitor oil exposure perturbed different sets of genes between F1 and F2 descendent embryos

The sets of genes that were perturbed by progenitor exposures were distinct in F1 and F2 descendants; there were no genes that were shared between the lists of genes showing a progenitor exposure effect in F1 and F2 embryos. F1 genes that are significantly perturbed by progenitor exposures (Figure 4A, pastel-fill squares) do not show a lineage exposure effect in F2 embryos (Figure 4A, bold-fill circles). The reciprocal is also apparent, where F2 genes that are significantly perturbed by progenitor exposures (Figure 4B, bold-fill circles) do not show a lineage exposure effect in F1 embryos. GO enrichment analyses also indicate little overlap in the biological functions inferred to be perturbed by progenitor exposures in F1 and F2 descendants (Figure 4C).

**Figure 4:**
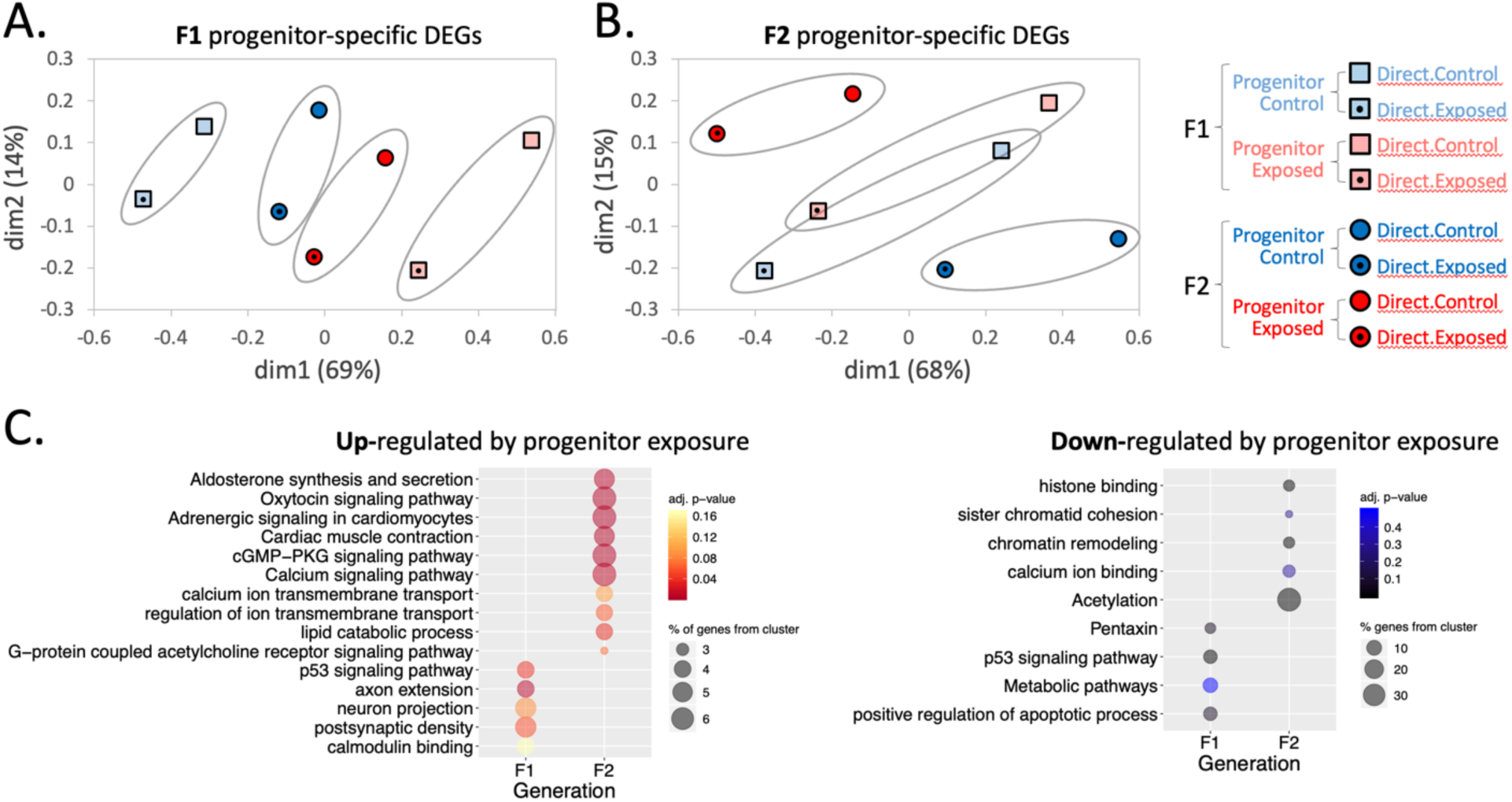
Progenitor exposure perturbs transcription during embryonic development in F1 and F2 descendants, but the sets of genes and molecular functions perturbed are different between F1 and F2 descendants. MDS plots include genes that are perturbed following progenitor exposure in F1 descendants (panel A) and in F2 descendants (panel B). Pastel squares and bold circles distinguish gene expression of F1 from F2 descendants, respectively. Red and blue fill distinguish gene expression of progenitor oil exposed from progenitor no-oil control lineages, respectively. Centered black dot or no dot distinguish gene expression of direct embryo oil exposed from embryo no-oil controls, respectively. For genes significantly perturbed by progenitor exposure in F1 descendants (panel A, pastel blue squares strongly distinguished from pastel red squares along dim1) we also show expression for those same genes in the F2 generation (panel A, bold blue and red circles). For genes significantly perturbed by progenitor exposure in F2 descendants (panel B, bold blue circles strongly distinguished from bold red circles along dim1) we also show expression for those same genes in the F1 generation (panel B, pastel blue and red squares). Genes that are perturbed by progenitor exposure in one generation of descendants tend to not be similarly perturbed by progenitor exposure in the other generation. Functional gene ontology (GO) enrichment analysis (panel C) was performed on genes that were up-regulated (left set in warm colors) or down-regulated (right set in cool colors) by progenitor exposures separately, and separately for genes perturbed in F1 and F2 descendants (four GO enrichment analyses). Since progenitor exposure perturbs the expression of different sets of genes in F1 and F2 descendants (panels A and B), and since the functional pathways perturbed by progenitor exposures in F1 and F2 descendants do not share GO terms (panel C), we conclude that progenitor exposures perturb different biological functions in F1 and F2 offspring.

### Transcriptomic responses to direct oil exposure in embryos were consistent across two generations and only subtly affected by progenitor exposure

There were many gene expression changes in response to direct embryonic oil exposure (Table 1). In late-stage embryos, we detected 639 DEGs and 716 DEGs in response to direct embryonic oil exposure in F1 and F2 embryos, respectively, regardless of their progenitor exposure lineage (Table 1). The transcriptional response to direct embryonic oiling was conserved between F1 and F2 embryos. Direct exposure-responsive genes in F1 embryos tended to show a conserved response in F2 embryos (Figure 5A). The reciprocal was also observed, where direct exposure-responsive genes in F2 embryos tended to show a conserved response in F1 embryos (Figure 5B). Gene ontology (GO) enrichment analyses for oil-responsive DEGs in late-stage F1 and F2 embryos show that many of the same pathways and biological functions are enriched in both sets of DEGs between generations (Figure 5C). These results are consistent with a conserved response to direct oiling between generations.

**Figure 5:**
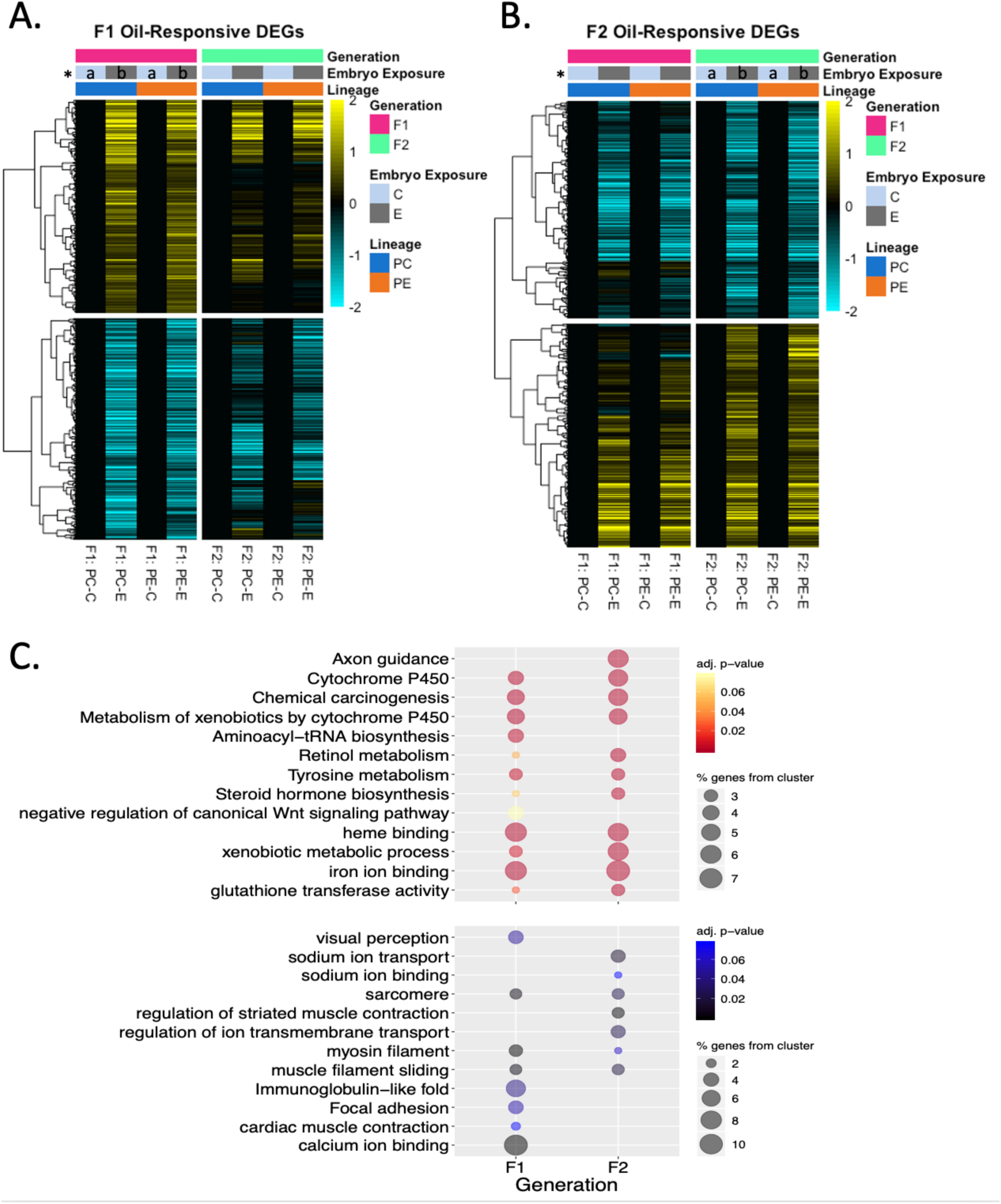
Direct embryonic oil exposure perturbs gene expression. The patterns of perturbation are consistent between progenitor exposed or control lineages and consistent between F1 and F2 embryos. Heatmaps include genes that are perturbed following direct embryonic exposure in F1 embryos (panel A) and in F2 embryos (panel B). Green and pink bars distinguish gene expression of F1 from F2 descendants. Light blue and grey bars distinguish gene expression of direct embryo oil exposed from embryo no-oil controls. Orange and dark blue bars distinguish gene expression of progenitor oil exposed from progenitor no-oil control lineages. For genes significantly perturbed by direct embryonic oil exposure in F1 embryos (panel A, left column highlighted in pink, where * a and b indicate treatments that have significant differences in gene expression) we also show expression for those same genes in F2 embryos (panel A, right column highlighted in green). For genes significantly perturbed by direct embryonic oil exposure in F2 embryos (panel B, right column highlighted in green, where * a and b indicate treatments that have significant differences in gene expression) we also show expression for those same genes in F1 embryos (panel B, left column highlighted in pink). Genes that are perturbed by direct embryonic oil exposure in one generation tend to be similarly perturbed in the other generation. Plotted are oil-responsive DEGs from F1 and F2 late-stage embryos after excluding DEGs with a significant main effect of parental exposure lineage or significant interaction between parental exposure lineage and direct embryo oil exposure treatment. Expression values are normalized to the control condition for each embryonic treatment groups to show relative expression changes in directly exposed versus no-oil control groups. Functional gene ontology (GO) enrichment analysis (panel C) was performed on genes that were up-regulated (top set in warm colors) or down-regulated (bottom set in cool colors) by direct embryonic exposures separately, and separately for genes perturbed in F1 and F2 embryos (four GO enrichment analyses). Since direct embryonic oil exposure tends to perturb the expression of the same sets of genes regardless of progenitor exposure, and tends to be conserved in F1 and F2 embryos (panels A and B), and since the functional pathways perturbed by direct embryonic exposures in F1 and F2 descendants largely overlap (panel C), we conclude that the molecular response to direct embryonic oil exposure is highly canalized and not influenced by exposures of progenitors.

Of the genes that showed a transcriptional response to direct embryonic oil exposure, very few varied in their response between progenitor exposure lineages. That is, there were very few genes that had a significant interaction between the direct embryonic exposure treatment and the progenitor exposure lineage treatment. Treatment interaction genes included 0 genes in early-stage embryos and 6 genes in late-stage embryos in the F1, and 4 genes and 53 genes in early-stage and late-stage embryos, respectively, in the F2 (Table 1). This suggests that the transcriptional response to embryonic direct oil exposure was only subtly influenced by progenitor exposure to oil, but more so in F2 descendants.

## DISCUSSION

Our results indicate that sub-lethal exposure to contaminating crude oil can cause biological impacts that propagate across at least two generations of descendant killifish. Specifically, we show that; 1) progenitor (F0) oil exposure alters morphological development, particularly in second-generation descendants (F2), and under some conditions can sensitize descendants to additional oiling, 2) progenitor oil exposure perturbs the transcriptome during embryonic development in F1 and F2 descendants, 3) the transcriptomic perturbation caused by progenitor exposure in F1 embryos is distinct from those observed in F2 embryos, 4) the transcriptomic response to direct embryonic oil exposure is not affected by progenitor oil exposure and is conserved between F1 and F2 generations. We conclude that progenitor oil exposures cause maladaptive outcomes in descendants (*purple* and *orange* outcomes in Figure 1). More specifically, the molecular data and some of the developmental morphology data indicate a general perturbation manifest in descendants that is independent of additional exposures (*orange* outcome in Figure 1), whereas a subset of the larval morphological data (interaction between direct oil exposure and lineage exposure for POL from the high acute direct oil concentration in F2) indicates sensitization of descendants to additional direct oil exposures (*purple* outcome in Figure 1).

The descendant effects that propagated from progenitor exposures were inconsistent between F1 and F2 generations. In F2 descendants, progenitor exposures caused a 3-4% reduction in larval length (Figure 3B, 3C). In F1 offspring, progenitor exposures caused a similar reduction in larval length, but this was not statistically significant (p = 0.09; Figure 3A). Exposures also perturbed pre-orbital length, where progenitor exposure sensitized F2 descendants to this outcome from direct oiling at the high dose (Figure 3F). A similar pattern was observed in F2 descendants exposed to the lower dose, and in F1 offspring, but these responses were not statistically significant (e.g., the greatest reductions in POL following direct embryo exposure was in embryos whose progenitors had been exposed; Figure 3E and 3D). Although impacts from direct exposures tended to be observed in both F1 and F2 larvae, lineage effects and interactions between lineage and direct exposure effects were most apparent in F2 descendants. Similarly, molecular responses to progenitor exposure varied across generations. Progenitor exposure perturbed the expression of more genes in F2 descendants than in F1 descendants; 40-fold more early-stage genes and nearly 2-fold more late-stage genes were perturbed by progenitor exposure in F2 compared to F1 descendants (Table 1). Furthermore, there was no overlap in the identity of genes and molecular pathways perturbed by progenitor exposure in F1 compared to F2 descendants (Figure 4).

Inconsistencies in the outcomes of progenitor exposures in different generations of descendants are likely a consequence of different mechanisms of across-generation transfer of information operating across different generations. In the first generation following exposure, multiple mechanisms may contribute (and interact in complex ways) to affect offspring biology. For example, females may load bioaccumulated chemicals into eggs such that developing embryos are effectively directly exposed (e.g., ^45^). Exposed females may also load exposure-perturbed molecules into eggs, such as RNA, proteins, and metabolites, which affect developmental programs in embryos ^46,47^. Exposures may also alter the energetic provisioning of eggs ^48^. These examples may be summarized as *maternal effects*. Furthermore, exposures may also cause heritable epigenetic changes in the genomes of eggs and sperm (genomic imprinting or “epimutations”, for example through histone modifications or DNA methylation), which may alter gene expression and development in early life ^11,12^. If there is no additional direct exposure throughout the F1 generation, then progenitor exposure effects that propagate into the F2 and beyond are likely mediated primarily by heritable epigenetic alterations ^49^. Consistent with this, we observed that *acetylation* and *histone binding* functions were enriched among the genes perturbed by progenitor exposure in F2 offspring (Figure 4C). Since many exposure-induced mechanisms contribute to F1 perturbations but only a subset contributes to F2 perturbations, then inconsistencies between F1 and F2 effects should be common. Indeed, this phenomenon is observed in many other transgenerational toxicology studies. For example, zebrafish progenitor exposure to dioxin caused greater transcriptome perturbations in F2 offspring compared to F1, with little overlap in the perturbed F1 and F2 genes ^50^. Similarly, PCB/PBDE exposures in zebrafish progenitors altered behavior and gene expression in F1-F4 offspring, with little overlap in the perturbed genes between descendent generations ^51^. Progenitor exposure to PAHs in sheepshead minnows caused reduced prey capture ability in only the F2 generation, and developmental abnormalities apparent in the F1 were diminished in the F2 ^27^. Progenitor exposure to PAH caused craniofacial malformations in F2 but not F1 descendants ^52^. Multigenerational effects of di-(2-ethylhexyl) phthalate (DEHP) exposure caused fewer socially investigative behaviors and increased exploratory behaviors in F1 mice, but these trends were reversed in F3 mice ^53^. Studies that track the transmission of exposure-induced chemicals, small molecules, energetic stores, and epigenetic alterations across generations, are necessary to illuminate the mechanisms underlying the complexity of transgenerational effects, and thereby increase our ability to predict and track health outcomes.

Progenitor exposures cause molecular perturbations in F1 and F2 descendants that are consistent with impacts on the morphology and physiology of descendants. Molecular responses also offer insights into the functional pathways leading to adverse outcomes. In a related transgenerational oil exposure study in *F. grandis* conducted by our group, we found that F1 and F2 descendants of oil-exposed progenitors had reduced swim performance (reduced mean critical swimming speed; *U*_crit_) ^54^. In the present study, genes that were perturbed in F2 late-stage embryos were enriched for those involved in heart function (*cardiac muscle contraction*, *adrenergic signaling in cardiomyocytes*, *calcium signaling pathway*, *calcium binding*; Figure 4C) which may be mechanistically linked to reduced swim performance later in life. Though direct exposure to oil is well-known to affect heart development ^55^ and swim performance in later-life ^56^, F2 fish with reduced swim performance had not been directly exposed to oil, and F1 fish may have been directly exposed to oil but only through any PAHs that may have been maternally loaded into eggs. These results suggest that impacts from progenitor oil exposures propagate to at least second-generation descendants through heritable perturbations of cellular memory (e.g., epigenetic imprinting), resulting in perturbation of cardiac development that diminishes later-life physiological performance. Although we also observed diminished swim performance that was similar between F1 as F2 descendants ^54^, we did not observe the same perturbation of cardiac gene expression in F1 as in F2 descendants (Figure 4C). It is plausible that perturbations of cellular memory were inherited in F1 descendants but did not manifest in the same gene expression perturbations as in F2 descendants because of complex interactions with maternal effects that masked the cardiac molecular response. Furthermore, we measured gene expression in whole embryos, where cardiac tissues constitute a small fraction; cardiac-specific epigenetic effects may have been masked by maternal effects that perturb gene expression across other tissues but become unmasked in F2 descendants in which maternal effects are of little influence. Follow-up studies using single-cell transcriptomics could expose perturbations that are specific to different cell types.

Progenitor oil exposure perturbed neurological gene expression in F1 descendant late-stage embryos (enriched GO functions included *axon extension*, *neuron projection*, *postsynaptic density*; Figure 4C). For F1 descendants we cannot distinguish the influence of maternal effects from heritable perturbations of cellular memory. Insofar as neuronal gene expression perturbations did not propagate into F2 descendants, we infer that maternal effects are more likely responsible than heritable epigenetic alterations. However, in medaka fish parental exposures to PAHs altered methylation and expression of neuronal genes in F1, which were associated with behavioral and other functional deficits ^57^. Direct exposures to crude oil have been shown to alter neurological gene expression ^58^, and cause behavioral deficits in fish ^59–61^. Furthermore, behavioral deficits have been detected in offspring of progenitors exposed to diverse toxicants including heavy metals ^62^, legacy pollutants ^51^, pharmaceuticals ^63^, endocrine-active compounds ^64^, hydrocarbons ^65^, and pesticides ^66^. We hypothesize that progenitor exposures to crude oil in killifish may cause behavioral impacts in at least first-generation offspring, perhaps similar to observations in sheepshead minnows ^27^ and medaka fish ^57^. Additional experiments are required to test this hypothesis.

The transcriptional responses to direct oil exposure during development were consistent between F1 and F2 embryos and were minimally affected by progenitor exposure. This canalized response to direct oiling involved the activation of molecular pathways that have clear mechanistic connections to adverse outcomes following oil exposures, and that are evolutionarily conserved across vertebrates ^55,67,68^. These mechanisms include activation of aryl hydrocarbon receptor signaling (e.g., reflected by enriched GO terms *cytochrome P450*, *xenobiotic metabolic process*, and *glutathione transferase activity*; Figure 5C), altered Wnt signaling (e.g., reflected by enriched GO term *negative regulation of canonical Wnt signaling pathway*; Figure 5C), and perturbation of intracellular calcium (e.g., reflected by enriched GO term *calcium ion binding*; Figure 5C), that manifest as cardiovascular system developmental deficits such as pericardial edema, malformed heart, altered heart rate, arrhythmia, impaired contractility, and reduced cardiac output ^55,69^. These results indicate that the molecular response to direct embryonic oil exposure was not altered (in an adaptive or maladaptive way) in lineages where progenitors had been exposed to oil.

The relatively small modifying effect of progenitor oil exposure on the transcriptional response to direct oiling (few genes with interactions between progenitor and direct exposures) is inconsistent with progenitor effects observed on the pre-orbital length (POL) response to direct oiling. POL is the distance between the snout and the anterior margin of the eye orbit. Changes in POL reflect craniofacial perturbations caused by PAH exposure in vertebrates (e.g., ^70–72)^. Descendants of oil-exposed adults were sensitized to this oil-induced effect, especially in F2 embryos exposed to the higher dose of oil (Figure 3F). We did not detect gene expression responses that provide insight into this morphological outcome; only 53 genes showed direct-exposure by progenitor-exposure interactions in their expression (Table 1). It is plausible that gene expression perturbations responsible for this morphological outcome occur in only a very small subset of embryonic tissues (e.g., as with dioxin-induced perturbation of FoxQ1b expression in the jaw primordium of developing zebrafish ^73^), such that they were not detectable in the multi-tissue context of whole embryos. It is also plausible that the molecular perturbations that underlie altered larval POL (measured immediately post-hatch) emerged later in development than when we assayed gene expression (at ∼15 days post-fertilization). Finally, although there was no functional enrichment within the 53 genes showing progenitor-by-direct oiling interactions, perhaps transcriptional changes of only a few of these genes are sufficient to cause craniofacial defects.

Our observation that progenitor exposure perturbs descendant biology, and that this perturbation manifests regardless of additional exposures, is consistent with other transgenerational toxicology literature (e.g., ^27,52,65,74^). Although some studies have indicated that progenitor exposures may sensitize or desensitize offspring to additional exposures, it is not always clear whether this is because of perturbations that are mechanistically associated with toxicant-specific mechanisms of toxicity or tolerance, or whether this reflects more generalized effects on health and performance. For example, one might imagine that descendants may be sensitized to additional exposures because of developmental malfunction or reduced energetic provisioning (e.g., ^48^), or desensitized because of inherited priming with metabolic or immune defense transcripts or proteins (e.g., ^7,26,46,75^) or through natural selection. We suggest that since toxicant exposures likely have no analog in evolutionary history, adaptive mechanisms of stressor-specific transgenerational transfer of information are unlikely, such that effects that span generations are likely to be general and negative. If they are general, then they may be difficult to predict and then detect. Therefore, comprehensive screening afforded by functional genomics or phenomics approaches may play important roles that serve to both generate and test hypotheses ^76,77^. Our results show that transgenerational outcomes following progenitor exposures are complex. Some outcomes may emerge independent of additional exposures, whereas other outcomes may only be revealed upon additional direct exposures. Similar complexity has been observed in other studies, where offspring from exposed parents were less sensitive to direct exposures, but offspring fitness was otherwise compromised in clean conditions ^7,78^. Furthermore, outcomes in one generation may be inconsistent with those that emerge in later generations. This is commonly observed and is likely the result of complex interactions between different combinations of mechanisms that are engaged to transmit transgenerational information across different generations. For example, chemical transfer, maternal effects, and genomic imprinting could interact to affect outcomes in F1 offspring, whereas F2 effects are more likely to be underpinned by genomic imprinting effects alone, or perhaps in combination with some lingering maternal effects. Additional complexity may emerge as a function of chemical novelty or as a function of dose and/or duration of exposure. Some pollutants are “natural”, insofar as they may have been commonly encountered in nature over the evolutionary history of wild species (e.g., some metals, some hydrocarbons), whereas other chemicals are less likely to have natural analogs (e.g., synthetic chemicals such as dioxins, PCBs, flame retardants, etc.). Toxicity of chemicals is a function of their dose, but for some chemicals dose-responses are not linear – that is, intermediate doses may be beneficial, whereas only high or low doses are toxic ^79^. For such chemicals, one might predict that transgenerational outcomes may be dependent on the dose or timing of exposure ^6,7^. Adaptive transgenerational outcomes are only plausible if the stimulus is a reliable indicator of environmental quality and the biotic responses has been shaped by evolutionary history ^5^, whereas exposures to novel agents are more likely to perturb information-transfer mechanisms and result in maladaptive outcomes. Since the chemical milieu experienced by diverse creatures in the Anthropocene includes novel agents encountered as discontinuous exposures, research into the importance of exposure time, and exposure timing coincident with susceptible life stages, will be important for enriching our ability to predict long-term outcomes of chemical exposures, thereby enhancing risk estimation.

## DATA AVAILABILITY

Raw sequence reads for transcriptomics are available at NCBI (BioProject PRJNA473014). Bioinformatics scripts for transcriptomics analyses are available as an R Markdown file (Appendix S2).

## ACKNOWLEDGEMENTS

Dr. Joanna Griffiths assisted with statistical analysis of morphometrics data. This research was supported by funding from the National Institutes of Environmental Health Sciences (1R01ES021934 to A.W. and F.G.) and from the National Science Foundation (OCE-1314567 to A.W. and F.G.).

## REFERENCES

(1) Bonduriansky, R.; Day, T. Nongenetic Inheritance and Its Evolutionary Implications. Annu. Rev. Ecol. Evol. Syst. 2009, 40 (1), 103–125. 10.1146/annurev.ecolsys.39.110707.173441.

(2) Jablonka, E.; Raz, G. Transgenerational Epigenetic Inheritance: Prevalence, Mechanisms, and Implications for the Study of Heredity and Evolution. Q Rev Biol 2009 2009, 131–176.

(3) Perez, M. F.; Lehner, B. Intergenerational and Transgenerational Epigenetic Inheritance in Animals. Nat. Cell Biol. 2019, 21 (2), 143–151. 10.1038/s41556-018-0242-9.

(4) Jablonka, E.; Oborny, B.; Molnár, I.; Kisdi, É.; Hofbauer, J.; Czárán, T. The Adaptive Advantage of Phenotypic Memory in Changing Environments. Philos. Trans. R. Soc. Lond. B. Biol. Sci. 1997, 350 (1332), 133–141. 10.1098/rstb.1995.0147.

(5) Donelan, S. C.; Hellmann, J. K.; Bell, A. M.; Luttbeg, B.; Orrock, J. L.; Sheriff, M. J.; Sih, A. Transgenerational Plasticity in Human-Altered Environments. Trends Ecol. Evol. 2020, 35 (2), 115–124. 10.1016/j.tree.2019.09.003.

(6) Ho, D. H.; Burggren, W. W. Parental Hypoxic Exposure Confers Offspring Hypoxia Resistance in Zebrafish (Danio Rerio). J. Exp. Biol. 2012, 215 (23), 4208–4216. 10.1242/jeb.074781.

(7) Bautista, N. M.; Burggren, W. W. Parental Stressor Exposure Simultaneously Conveys Both Adaptive and Maladaptive Larval Phenotypes through Epigenetic Inheritance in the Zebrafish (Danio Rerio). J. Exp. Biol. 2019, 222 (17), jeb208918. 10.1242/jeb.208918.

(8) Reed, T. E.; Waples, R. S.; Schindler, D. E.; Hard, J. J.; Kinnison, M. T. Phenotypic Plasticity and Population Viability: The Importance of Environmental Predictability. Proc. R. Soc. B Biol. Sci. 2010, 277 (1699), 3391–3400. 10.1098/rspb.2010.0771.

(9) Agrawal, A. A.; Laforsch, C.; Tollrian, R. Transgenerational Induction of Defences in Animals and Plants. Nature 1999, 401 (6748), 60–63. 10.1038/43425.

(10) Salinas, S.; Munch, S. B. Thermal Legacies: Transgenerational Effects of Temperature on Growth in a Vertebrate. Ecol. Lett. 2012, 15 (2), 159–163. 10.1111/j.1461-0248.2011.01721.x.

(11) Szyf, M. The Dynamic Epigenome and Its Implications in Toxicology. Toxicol. Sci. 2007, 100 (1), 7–23. 10.1093/toxsci/kfm177.

(12) Nilsson, E. E.; Ben Maamar, M.; Skinner, M. K. Role of Epigenetic Transgenerational Inheritance in Generational Toxicology. Environ. Epigenetics 2022, 8 (1), dvac001. 10.1093/eep/dvac001.

(13) Rozas, L. P.; Reed, D. J. Nekton Use of Marsh-Surface Habitats in Louisiana (USA) Deltaic Salt Marshes Undergoing Submergence. Mar. Ecol. Prog. Ser. 1993, 96 (2), 147–157. Doi 10.3354/Meps096147.

(14) Rozas, L. P.; Zimmerman, R. J. Small-Scale Patterns of Nekton Use among Marsh and Adjacent Shallow Nonvegetated Areas of the Galveston Bay Estuary, Texas (USA). Mar. Ecol.-Prog. Ser. 2000, 193, 217–239.

(15) Jernelöv, A. The Threats from Oil Spills: Now, Then, and in the Future. Ambio 2010, 39 (5– 6), 353–366. 10.1007/s13280-010-0085-5.

(16) Incardona, J. P. Molecular Mechanisms of Crude Oil Developmental Toxicity in Fish. Arch. Environ. Contam. Toxicol. 2017, 73 (1), 19–32. 10.1007/s00244-017-0381-1.

(17) Pasparakis, C.; Esbaugh, A. J.; Burggren, W.; Grosell, M. Physiological Impacts of Deepwater Horizon Oil on Fish. Comp. Biochem. Physiol. Part C Toxicol. Pharmacol. 2019, 224, 108558. 10.1016/j.cbpc.2019.06.002.

(18) Grosell, M.; Pasparakis, C. Physiological Responses of Fish to Oil Spills. Annu. Rev. Mar. Sci. 2021, 13 (1), 137–160. 10.1146/annurev-marine-040120-094802.

(19) Hansen, B. H.; Tarrant, A. M.; Salaberria, I.; Altin, D.; Nordtug, T.; Overjordet, I. B. Maternal Polycyclic Aromatic Hydrocarbon (PAH) Transfer and Effects on Offspring of Copepods Exposed to Dispersed Oil with and without Oil Droplets. J. Toxicol. Environ. Health-Part - Curr. Issues 2017, 80 (16–18), 881–894. 10.1080/15287394.2017.1352190.

(20) Yang, J.; Chatterjee, N.; Kim, Y.; Roh, J.-Y.; Kwon, J.-H.; Park, M.-S.; Choi, J. Histone Methylation-Associated Transgenerational Inheritance of Reproductive Defects in Caenorhabditis Elegans Exposed to Crude Oil under Various Exposure Scenarios. Chemosphere 2018, 200, 358–365. 10.1016/j.chemosphere.2018.02.080.

(21) Duan, M.; Xiong, D.; Bai, X.; Gao, Y.; Xiong, Y.; Gao, X.; Ding, G. Transgenerational Effects of Heavy Fuel Oil on the Sea Urchin Strongylocentrotus Intermedius Considering Oxidative Stress Biomarkers. Mar. Environ. Res. 2018, 141, 138–147. 10.1016/j.marenvres.2018.08.010.

(22) Nikinmaa, M.; Suominen, E.; Anttila, K. Water-Soluble Fraction of Crude Oil Affects Variability and Has Transgenerational Effects in Daphnia Magna. Aquat. Toxicol. 2019, 211, 137–140. 10.1016/j.aquatox.2019.04.004.

(23) Sun, L.; Zuo, Z.; Chen, M.; Chen, Y.; Wang, C. Reproductive and Transgenerational Toxicities of Phenanthrene on Female Marine Medaka (Oryzias Melastigma). Aquat. Toxicol. 2015, 162, 109–116. 10.1016/j.aquatox.2015.03.013.

(24) Wang, Y.; Zhong, H.; Wang, C.; Gao, D.; Zhou, Y.; Zuo, Z. Maternal Exposure to the Water Soluble Fraction of Crude Oil, Lead and Their Mixture Induces Autism-like Behavioral Deficits in Zebrafish (Danio Rerio) Larvae. Ecotoxicol. Environ. Saf. 2016, 134, 23–30. 10.1016/j.ecoenv.2016.08.009.

(25) Philibert, D. A.; Lyons, D. D.; Qin, R.; Huang, R.; El-Din, M. G.; Tierney, K. B. Persistent and Transgenerational Effects of Raw and Ozonated Oil Sands Process-Affected Water Exposure on a Model Vertebrate, the Zebrafish. Sci. Total Environ. 2019, 693, 133611. 10.1016/j.scitotenv.2019.133611.

(26) Bautista, N. M.; Crespel, A.; Crossley, J.; Padilla, P.; Burggren, W. Parental Transgenerational Epigenetic Inheritance Related to Dietary Crude Oil Exposure in Danio Rerio. J. Exp. Biol. 2020, 223 (16), jeb222224. 10.1242/jeb.222224.

(27) Jasperse, L.; Levin, M.; Rogers, K.; Perkins, C.; Bosker, T.; Griffitt, R. J.; Sepúlveda, M. S.; De Guise, S. Transgenerational Effects of Polycyclic Aromatic Hydrocarbon Exposure on Sheepshead Minnows (Cyprinodon Variegatus). Environ. Toxicol. Chem. 2019, 38 (3), 638–649. 10.1002/etc.4340.

(28) Carls, M. G.; Hose, J. E.; Thomas, R. E.; Rice, S. D. Exposure of Pacific Herring to Weathered Crude Oil: Assessing Effects on Ova. Environ. Toxicol. Chem. 2000, 19 (6), 1649–1659. 10.1002/etc.5620190624.

(29) Kennedy, C. J.; Farrell, A. P. Effects of Exposure to the Water-Soluble Fraction of Crude Oil on the Swimming Performance and the Metabolic and Ionic Recovery Post-Exercise in Pacific Herring (Clupea Pallasi). Environ. Toxicol. Chem. 2006, 25 (10), 2715–2724. 10.1897/05-504R.1.

(30) Pilcher, W.; Miles, S.; Tang, S.; Mayer, G.; Whitehead, A. Genomic and Genotoxic Responses to Controlled Weathered-Oil Exposures Confirm and Extend Field Studies on Impacts of the Deepwater Horizon Oil Spill on Native Killifish. PLoS ONE 2014, 9 (9), e106351. Artn E106351 10.1371/Journal.Pone.0106351.

(31) Incardona, J. P.; Swarts, T. L.; Edmunds, R. C.; Linbo, T. L.; Aquilina-Beck, A.; Sloan, C. A.; Gardner, L. D.; Block, B. A.; Scholz, N. L. Exxon Valdez to Deepwater Horizon: Comparable Toxicity of Both Crude Oils to Fish Early Life Stages. Aquat. Toxicol. 2013, 142–143, 303–316. 10.1016/j.aquatox.2013.08.011.

(32) Fox, J.; Weisberg, S. An R Companion to Applied Regression, Third.; Sage: Thousand Oaks CA, 2019.

(33) Lenth, R. V. Emmeans: Estimated Marginal Means, Aka Least-Squares Means; 2023.

(34) Posit team. RStudio: Integrated Development Environment for R; Posit Software, PBC: Boston, MA, 2023.

(35) Andrew, S. FastQC: A Quality Control Tool for High Throughput Sequence Data.; 2010.

(36) Bolger, A. M.; Lohse, M.; Usadel, B. Trimmomatic: A Flexible Read Trimming Tool for Illumina NGS Data; 2014.

(37) Patro, R.; Duggal, G.; Love, M. I.; Irizarry, R. A.; Kingsford, C. Salmon Provides Fast and Bias-Aware Quantification of Transcript Expression. Nat. Methods 2017, 14, 417.

(38) Reid, N. M.; Jackson, C. E.; Gilbert, D.; Minx, P.; Montague, M. J.; Hampton, T. H.; Helfrich, L. W.; King, B. L.; Nacci, D. E.; Aluru, N.; Karchner, S. I.; Colbourne, J. K.; Hahn, M. E.; Shaw, J. R.; Oleksiak, M. F.; Crawford, D. L.; Warren, W. C.; Whitehead, A. The Landscape of Extreme Genomic Variation in the Highly Adaptable Atlantic Killifish. Genome Biol. Evol. 2017, 9 (3), 659–676. 10.1093/gbe/evx023.

(39) Soneson, C.; Love, M. I.; Robinson, M. D. Differential Analyses for RNA-Seq: Transcript-Level Estimates Improve Gene-Level Inferences. F1000Research 2016, 4, 1521. 10.12688/f1000research.7563.2.

(40) Bozinovic, G.; Oleksiak, M. F. Embryonic Gene Expression among Pollutant Resistant and Sensitive Fundulus Heteroclitus Populations. Aquat. Toxicol. 2010, 98 (3), 221–229. 10.1016/j.aquatox.2010.02.022.

(41) White, R. J.; Collins, J. E.; Sealy, I. M.; Wali, N.; Dooley, C. M.; Digby, Z.; Stemple, D. L.; Murphy, D. N.; Billis, K.; Hourlier, T.; Füllgrabe, A.; Davis, M. P.; Enright, A. J.; Busch-Nentwich, E. M. A High-Resolution mRNA Expression Time Course of Embryonic Development in Zebrafish. eLife 2017, 6, e30860. 10.7554/eLife.30860.

(42) Briggs, J. A.; Weinreb, C.; Wagner, D. E.; Megason, S.; Peshkin, L.; Kirschner, M. W.; Klein, A. M. The Dynamics of Gene Expression in Vertebrate Embryogenesis at Single-Cell Resolution. Science 2018, 360 (6392), eaar5780. 10.1126/science.aar5780.

(43) R Development Core Team, R. R: A Language and Environment for Statistical Computing; Team, R. D. C., Ed.; R Foundation for Statistical Computing; R Foundation for Statistical Computing, 2011; Vol. 1. 10.1007/978-3-540-74686-7.

(44) Huang, D. W.; Sherman, B. T.; Lempicki, R. A. Systematic and Integrative Analysis of Large Gene Lists Using DAVID Bioinformatics Resources. Nat. Protoc. 2009, 4 (1), 44–57. 10.1038/nprot.2008.211.

(45) Martins, M. F.; Costa, P. G.; Bianchini, A. Maternal Transfer of Polycyclic Aromatic Hydrocarbons in an Endangered Elasmobranch, the Brazilian Guitarfish. Chemosphere 2021, 263, 128275. 10.1016/j.chemosphere.2020.128275.

(46) Hasselquist, D.; Nilsson, J.-Å. Maternal Transfer of Antibodies in Vertebrates: Trans-Generational Effects on Offspring Immunity. Philos. Trans. R. Soc. B Biol. Sci. 2008, 364 (1513), 51–60. 10.1098/rstb.2008.0137.

(47) Klosin, A.; Lehner, B. Mechanisms, Timescales and Principles of Trans-Generational Epigenetic Inheritance in Animals. Curr. Opin. Genet. Dev. 2016, 36, 41–49. 10.1016/j.gde.2016.04.001.

(48) Kozal, J. S.; Jayasundara, N.; Massarsky, A.; Lindberg, C. D.; Oliveri, A. N.; Cooper, E. M.; Levin, E. D.; Meyer, J. N.; Giulio, R. T. D. Mitochondrial Dysfunction and Oxidative Stress Contribute to Cross-Generational Toxicity of Benzo(a)Pyrene in Danio Rerio. Aquat. Toxicol. 2023, 263, 106658. 10.1016/j.aquatox.2023.106658.

(49) Youngson, N. A.; Whitelaw, E. Transgenerational Epigenetic Effects. Annu. Rev. Genomics Hum. Genet. 2008, 9 (1), 233–257. 10.1146/annurev.genom.9.081307.164445.

(50) Meyer, D. N.; Baker, B. B.; Baker, T. R. Ancestral TCDD Exposure Induces Multigenerational Histologic and Transcriptomic Alterations in Gonads of Male Zebrafish. Toxicol. Sci. 2018, 164 (2), 603–612. 10.1093/toxsci/kfy115.

(51) Alfonso, S.; Blanc, M.; Joassard, L.; Keiter, S. H.; Munschy, C.; Loizeau, V.; Bégout, M.-L.; Cousin, X. Examining Multi-and Transgenerational Behavioral and Molecular Alterations Resulting from Parental Exposure to an Environmental PCB and PBDE Mixture. Aquat. Toxicol. 2019, 208, 29–38. 10.1016/j.aquatox.2018.12.021.

(52) Corrales, J.; Thornton, C.; White, M.; Willett, K. L. Multigenerational Effects of Benzo[a]Pyrene Exposure on Survival and Developmental Deformities in Zebrafish Larvae. Aquat. Toxicol. 2014, 148, 16–26. 10.1016/j.aquatox.2013.12.028.

(53) Quinnies, K. M.; Harris, E. P.; Snyder, R. W.; Sumner, S. S.; Rissman, E. F. Direct and Transgenerational Effects of Low Doses of Perinatal Di-(2-Ethylhexyl) Phthalate (DEHP) on Social Behaviors in Mice. PLOS ONE 2017, 12 (2), e0171977. 10.1371/journal.pone.0171977.

(54) Hess, C.; Little, L.; Brown, C.; Kaller, M.; Galvez, F. Transgenerational Effects of Parental Crude Oil Exposure on the Morphology of Adult Fundulus Grandis. Aquat. Toxicol. 2022, 249, 106209. 10.1016/j.aquatox.2022.106209.

(55) Cherr, G. N.; Fairbairn, E.; Whitehead, A. Impacts of Petroleum-Derived Pollutants on Fish Development. Annu. Rev. Anim. Biosci. Vol 5 2017, 5, 185–203. 10.1146/annurev-animal-022516-022928.

(56) Hicken, C. E.; Linbo, T. L.; Baldwin, D. H.; Willis, M. L.; Myers, M. S.; Holland, L.; Larsen, M.; Stekoll, M. S.; Rice, S. D.; Collier, T. K.; Scholz, N. L.; Incardona, J. P. Sublethal Exposure to Crude Oil during Embryonic Development Alters Cardiac Morphology and Reduces Aerobic Capacity in Adult Fish. Proc. Natl. Acad. Sci. 2011, 108 (17), 7086–7090. 10.1073/pnas.1019031108.

(57) Wan, T.; Au, D. W.-T.; Mo, J.; Chen, L.; Cheung, K.-M.; Kong, R. Y.-C.; Seemann, F. Assessment of Parental Benzo[a]Pyrene Exposure-Induced Cross-Generational Neurotoxicity and Changes in Offspring Sperm DNA Methylome in Medaka Fish. Environ. Epigenetics 2022, 8 (1), dvac013. 10.1093/eep/dvac013.

(58) Xu, E. G.; Khursigara, A. J.; Li, S.; Esbaugh, A. J.; Dasgupta, S.; Volz, D. C.; Schlenk, D. mRNA-miRNA-Seq Reveals Neuro-Cardio Mechanisms of Crude Oil Toxicity in Red Drum (Sciaenops Ocellatus). Environ. Sci. Technol. 2019, 53 (6), 3296–3305. 10.1021/acs.est.9b00150.

(59) Bautista, N. M.; Pothini, T.; Meng, K.; Burggren, W. W. Behavioral Consequences of Dietary Exposure to Crude Oil Extracts in the Siamese Fighting Fish (Betta Splendens). Aquat. Toxicol. 2019, 207, 34–42. 10.1016/j.aquatox.2018.11.025.

(60) Rowsey, L. E.; Johansen, J. L.; Khursigara, A. J.; Esbaugh, A. J. Oil Exposure Impairs Predator–Prey Dynamics in Larval Red Drum (Sciaenops Ocellatus). Mar. Freshw. Res. 2019, 71 (1), 99–106. 10.1071/MF18263.

(61) Armstrong, T.; Khursigara, A. J.; Killen, S. S.; Fearnley, H.; Parsons, K. J.; Esbaugh, A. J. Oil Exposure Alters Social Group Cohesion in Fish. Sci. Rep. 2019, 9 (1), 13520. 10.1038/s41598-019-49994-1.

(62) Carvan, M. J.; Kalluvila, T. A.; Klingler, R. H.; Larson, J. K.; Pickens, M.; Mora-Zamorano, F. X.; Connaughton, V. P.; Sadler-Riggleman, I.; Beck, D.; Skinner, M. K. Mercury-Induced Epigenetic Transgenerational Inheritance of Abnormal Neurobehavior Is Correlated with Sperm Epimutations in Zebrafish. PLOS ONE 2017, 12 (5), e0176155. 10.1371/journal.pone.0176155.

(63) De Serrano, A. R.; Hughes, K. A.; Rodd, F. H. Paternal Exposure to a Common Pharmaceutical (Ritalin) Has Transgenerational Effects on the Behaviour of Trinidadian Guppies. Sci. Rep. 2021, 11 (1), 3985. 10.1038/s41598-021-83448-x.

(64) Volkova, K.; Reyhanian Caspillo, N.; Porseryd, T.; Hallgren, S.; Dinnetz, P.; Olsén, H.; Porsch Hällström, I. Transgenerational Effects of 17α-Ethinyl Estradiol on Anxiety Behavior in the Guppy, Poecilia Reticulata. Gen. Comp. Endocrinol. 2015, 223, 66–72. 10.1016/j.ygcen.2015.09.027.

(65) Knecht, A. L.; Truong, L.; Marvel, S. W.; Reif, D. M.; Garcia, A.; Lu, C.; Simonich, M. T.; Teeguarden, J. G.; Tanguay, R. L. Transgenerational Inheritance of Neurobehavioral and Physiological Deficits from Developmental Exposure to Benzo[a]Pyrene in Zebrafish. Toxicol. Appl. Pharmacol. 2017, 329, 148–157. 10.1016/j.taap.2017.05.033.

(66) Pompermaier, A.; Tamagno, W. A.; Alves, C.; Barcellos, L. J. G. Persistent and Transgenerational Effects of Pesticide Residues in Zebrafish. Comp. Biochem. Physiol. Part C Toxicol. Pharmacol. 2022, 262, 109461. 10.1016/j.cbpc.2022.109461.

(67) Marris, C. R.; Kompella, S. N.; Miller, M. R.; Incardona, J. P.; Brette, F.; Hancox, J. C.; Sørhus, E.; Shiels, H. A. Polyaromatic Hydrocarbons in Pollution: A Heart-Breaking Matter. J. Physiol. 2020, 598 (2), 227–247. 10.1113/JP278885.

(68) Takeshita, R.; Bursian, S. J.; Colegrove, K. M.; Collier, T. K.; Deak, K.; Dean, K. M.; De Guise, S.; DiPinto, L. M.; Elferink, C. J.; Esbaugh, A. J.; Griffitt, R. J.; Grosell, M.; Harr, K. E.; Incardona, J. P.; Kwok, R. K.; Lipton, J.; Mitchelmore, C. L.; Morris, J. M.; Peters, E. S.; Roberts, A. P.; Rowles, T. K.; Rusiecki, J. A.; Schwacke, L. H.; Smith, C. R.; Wetzel, D. L.; Ziccardi, M. H.; Hall, A. J. A Review of the Toxicology of Oil in Vertebrates: What We Have Learned Following the Deepwater Horizon Oil Spill. J. Toxicol. Environ. Health Part B 2021, 24 (8), 355–394. 10.1080/10937404.2021.1975182.

(69) Sorhus, E.; Incardona, J. P.; Furmanek, T.; Goetz, G. W.; Scholz, N. L.; Meier, S.; Edvardsen, R. B.; Jentoft, S. Novel Adverse Outcome Pathways Revealed by Chemical Genetics in a Developing Marine Fish. Elife 2017, 6, e20707. ARTN e20707 10.7554/eLife.20707.

(70) Incardona, J. P.; Collier, T. K.; Scholz, N. L. Defects in Cardiac Function Precede Morphological Abnormalities in Fish Embryos Exposed to Polycyclic Aromatic Hydrocarbons. Toxicol. Appl. Pharmacol. 2004, 196 (2), 191–205. DOI 10.1016/j.taap.2003.11.026.

(71) Sørhus, E.; Incardona, J. P.; Karlsen, Ø.; Linbo, T.; Sørensen, L.; Nordtug, T.; van der Meeren, T.; Thorsen, A.; Thorbjørnsen, M.; Jentoft, S.; Edvardsen, R. B.; Meier, S. Crude Oil Exposures Reveal Roles for Intracellular Calcium Cycling in Haddock Craniofacial and Cardiac Development. Sci. Rep. 2016, 6 (1), 31058. 10.1038/srep31058.

(72) Sørhus, E.; Nakken, C. L.; Donald, C. E.; Ripley, D. M.; Shiels, H. A.; Meier, S. Cardiac Toxicity of Phenanthrene Depends on Developmental Stage in Atlantic Cod (*Gadus Morhua*). Sci. Total Environ. 2023, 881, 163484. 10.1016/j.scitotenv.2023.163484.

(73) Planchart, A.; Mattingly, C. J. 2,3,7,8-Tetrachlorodibenzo-p-Dioxin Upregulates FoxQ1b in Zebrafish Jaw Primordium. Chem. Res. Toxicol. 2010, 23 (3), 480–487. 10.1021/tx9003165.

(74) Crews, D.; Gillette, R.; Scarpino, S. V.; Manikkam, M.; Savenkova, M. I.; Skinner, M. K. Epigenetic Transgenerational Inheritance of Altered Stress Responses. Proc. Natl. Acad. Sci. 2012, 109 (23), 9143–9148. 10.1073/pnas.1118514109.

(75) Tsui, M. T. K.; Wang, W.-X. Multigenerational Acclimation of Daphnia Magna to Mercury: Relationships between Biokinetics and Toxicity. Environ. Toxicol. Chem. 2005, 24 (11), 2927–2933. 10.1897/05-085R.1.

(76) Cheng, K. C.; Katz, S. R.; Lin, A. Y.; Xin, X.; Ding, Y. Whole-Organism Cellular Pathology: A Systems Approach to Phenomics. In Advances in Genetics; Elsevier, 2016; Vol. 95, pp 89–115. 10.1016/bs.adgen.2016.05.003.

(77) Trego, M. L.; Brown, C. A.; Dubansky, B.; Hess, C. D.; Galvez, F.; Whitehead, A. Transcriptome Profiling in Conservation Physiology: Mechanistic Insights into Organism-Environment Interactions to Both Test and Generate Hypotheses. In Conservation Physiology: Applications for Wildlife Conservation and Management; OXFORD UNIV PRESS: S.l., 2020.

(78) Marshall, D. J. Transgenerational Plasticity in the Sea: Context-Dependent Maternal Effects Across the Life History. Ecology 2008, 89 (2), 418–427. 10.1890/07-0449.1.

(79) Calabrese, E. J.; Baldwin, L. A. Hormesis: The Dose-Response Revolution. Annu. Rev. Pharmacol. Toxicol. 2003, 43, 175–197. 10.1146/annurev.pharmtox.43.100901.140223.

