## Appendix S1 for "Adult killifish exposure to crude oil perturbs embryonic gene expression and larval morphology in first- and second-generation offspring"

**Appendix S1.** List of the 39 polycyclic aromatic compounds, the classification of the PAC (Low molecular weight, LMH or high molecular weight, HMW), the number of ring structures and the carbon numbers.

| **Analyte** | **PAC Type Classification** | **Carbon No.** |
| --- | --- | --- |
| Naphthalene | LMW: 2-ring | 10 |
| C2-Naphthalenes | LMW: 2-ring | 12 |
| C3-Naphthalenes | LMW: 2-ring | 13 |
| C4-Naphthalenes | LMW: 2-ring | 14 |
| Acenaphthene | LMW: 3-ring | 12 |
| Acenaphthylene | LMW: 3-ring | 12 |
| Biphenyl | LMW: 2-ring | 12 |
| Dibenzothiophene | LMW: 3-ring | 12 |
| C1-Dibenzothiophenes | LMW: 3-ring | 13 |
| C2-Dibenzothiophenes | LMW: 3-ring | 14 |
| C3-Dibenzothiophenes | LMW: 3-ring | 15 |
| C4-Dibenzothiophenes | LMW: 3-ring | 16 |
| Fluorene | LMW: 3-ring | 13 |
| C1-Fluorenes | LMW: 3-ring | 14 |
| C2-Fluorenes | LMW: 3-ring | 15 |
| C3-Fluorenes | LMW: 3-ring | 16 |
| Anthracene | LMW: 3-ring | 14 |
| Phenanthrene | LMW: 3-ring | 14 |
| C1-Phenanthrenes/Anthracenes | LMW: 3-ring | 15 |
| C2-Phenanthrenes/Anthracenes | LMW: 3-ring | 16 |
| C3-Phenanthrenes/Anthracenes | LMW: 3-ring | 17 |
| C4-Phenanthrenes/Anthracenes | LMW: 3-ring | 18 |
| Fluoranthene | HMW: 4-ring | 16 |
| Pyrene | HMW: 4-ring | 16 |
| C1-Fluoranthenes/Pyrenes | HMW: 4-ring | 17 |
| C2-Fluoranthenes/Pyrenes | HMW: 4-ring | 18 |
| C3-Fluoranthenes/Pyrenes | HMW: 4-ring | 19 |
| C4-Fluoranthenes/Pyrenes | HMW: 4-ring | 20 |
| Chrysene | HMW: 4-ring | 18 |
| C1-Chrysenes | HMW: 4-ring | 19 |
| C2-Chrysenes | HMW: 4-ring | 20 |
| C3-Chrysenes | HMW: 4-ring | 21 |
| C4-Chrysenes | HMW: 4-ring | 22 |
| Benzo(b)fluoranthene | HMW: 5-ring | 20 |
| Benzo(a)pyrene | HMW: 5-ring | 20 |
| Benzo(e)pyrene | HMW: 5-ring | 20 |
| Perylene | HMW: 5-ring | 20 |
| Benzo(g,h,i)perylene | HMW: 6-ring | 22 |
| Indeno(1,2,3-cd)pyrene | HMW: 6-ring | 22 |
